# A self without a “we”: Individuals with Borderline personality disorder show signatures of disrupted social basis functions

**DOI:** 10.64898/2026.09.21.752665

**Authors:** Marco K. Wittmann, Yongling Lin, Simon Ciranka, Sabrina Mittermeier, Marcel Romanos, Anthony S. David, Arne Bürger, Klara Gregorova, Andrea M.F. Reiter

**Affiliations:** Department of Experimental Psychology, University College London, 26 Bedford Way, London WC1H 0AP, UK; Max Planck UCL Centre for Computational Psychiatry and Ageing Research, University College London, Russell Square House 10-12 Russell Square, London WC1B 5EH, UK; Center for Adaptive Rationality, Max Planck Institute for Human Development, Lentzeallee 94, 14195 Berlin, Germany; Department of Child and Adolescent Psychiatry, Psychosomatics and Psychotherapy, University Hospital Würzburg, Germany; German Center of Prevention Research on Mental Health, Würzburg, Germany; Division of Psychiatry, University College London, London, UK; Department of Psychology, Julius-Maximilians-University of Würzburg, Würzburg, Germany

**Author notes:** Contributed equally (Marco K. Wittmann, Yongling Lin).

## Abstract

A capacity to navigate social relationships is fundamental to human wellbeing. However, the cognitive mechanisms underlying social dysfunction in psychiatric conditions remain poorly understood. Using an abstract multi-person decision making task that isolates the relational structure of multi-person interactions, we compare non-clinical controls (NCC, n=51) and individuals with BPD symptoms (n=46), a condition characterized by disturbances in self–other boundaries and chronic feelings of loneliness. Generally, we found a pattern of less self-centered decision-making in BPD. More specifically, individuals with BPD showed signatures of perceiving opponents as a coherent group (“them”), but a markedly reduced tendency to represent themselves as part of their own team (“us”). This pattern of results points towards a disruption of a primary social basis function. Our laboratory findings extend to experienced loneliness in daily life as measured by ecological momentary assessments (EMA). Together, this suggests that BPD is characterized by an unbalanced representation of social structure, with an attenuated “we” alongside a heightened “they”.

## Introduction

A capacity to navigate social environments with flexibility is fundamental to human functioning and wellbeing. Dysfunctions therein cause significant distress and impairment in everyday life. Yet, flexible social behavior is no trivial task. It requires the continuous integration of self-related and other-related information. People must understand how they are embedded in social structures, characterized by relationships of varying valence, ranging from mutually beneficial cooperation to competition (1). Our recent work has established a mechanistic account of how humans are able to master this complex task, using an abstract multi-person decision-making task that strips away the naturalistic features of real social interactions while preserving their underlying relational structure (2–4): We showed that humans rely on social building blocks that, in combination, can produce different interaction patterns and that are then linked to social identities such as “me” and “others”.

Disturbances in both self-concept and relationships with others are core features of individuals with borderline personality disorder (BPD). Indeed, there is a longstanding clinical tradition of conceptualizing BPD as characterized by disturbances in self–other boundaries, i.e. in the integration of self- and other-related information (5–8). For example, a person with BPD might say “I do not know who I am”, reflecting an unstable sense of self, and alongside both fear of being dependent on others and being left alone (9). Problems with self-other boundaries in BPD have been associated with greater symptom severity and less favorable treatment outcomes (10,11). They are also thought to underpin interpersonal problems in BPD (12). Individuals with BPD experience chronic loneliness, even in the absence of social isolation, compared to both non-clinical controls and even those with other psychiatric disorders (13–16), which is currently not targeted by first-line psychotherapeutic treatments (17,18). In sum, individuals with BPD are known to experience disturbances in self-other boundaries and an increased susceptibility to chronic loneliness. However, despite these longstanding clinical observations, we have limited knowledge about the cognitive and computational mechanisms underlying altered self-other interactions and how they relate to loneliness in BPD.

Recent experimental studies have yielded seemingly contradictory findings in BPD. On the one hand, BPD patients appear less influenced by behaviors observed in others and resist updating beliefs about themselves based on feedback from others (19,20), pointing to a self that might be more shielded from external influences. On the other hand, BPD patients can be more prone to incorporating other-generated actions and perceptions into the self (21), and to conflating mental states of the self and other (22,23). Rather than reflecting a simple deficit in self–other distinction, these findings together might suggest that individuals with BPD struggle with *flexibly regulating* a boundary between self and other (24).

Yet understanding *when and why* these shifts occur requires studying self–other boundaries within a complex multi-person relational context, while stripping away other naturalistic features of social interactions that make it hard to quantify the underlying computations. We focus on a simplified model of the relationships between multiple agents, as such relationships are at the heart of everyday life experiences and require individuals to flexibly integrate information about oneself, others, and their interrelationships simultaneously. Previous studies on social decision-making in BPD have primarily focused on dyadic scenarios (19,23,25). Further, laboratory studies of social cognition and observational studies of loneliness in BPD have largely remained disconnected: the former offer mechanistic precision but limited ecological validity, while the latter capture real-world experience but lack a formal account of the underlying processes. As a result, the hypothesis that disruptions in self–other distinction directly contribute to the pervasive social dysfunction experienced by individuals with BPD in daily life (12) has remained hard to test.

To address these limitations, we draw on recent findings in healthy adults showing that the relational structure underlying social interaction can be represented through fundamental mathematical constructs – known as basis functions – that can be combined to flexibly encode and compress social information (4). The key intuition is this: rather than representing every possible social relationship from scratch, complex interaction patterns can be decomposed into a small set of reusable building blocks. Much like how a limited palette of colors can be mixed to produce any shade, combinations of these basis functions can flexibly represent a wide range of relational dynamics (26,27). For example, we have shown that people encode the contrast between their own group and an outgroup (“us vs. them”) as a primary social basis function (2,4). Crucially, disruptions in these basis functions can produce failures of self– other distinction (28–31), making them a promising mechanistic account of the social difficulties observed in BPD.

Here, to study how individuals with BPD differ in their use of social basis functions, and how this relates to everyday social dysfunction, we compare behavior of adolescents and young adults with BPD symptoms (n=46; Table 1) and non-clinical controls (NCC, n=51) using an abstract multi-person social decision-making task (2–4), in which participants were assigned different teams and had to judge if their own or the other team was better. Subsequently, we relate this to ecological momentary assessment ratings of loneliness in the same cohorts. First, we replicate our previous observation that NCCs exhibit more efficient social information weighting for themselves compared to for a partner. This effect is strikingly absent in individuals with BPD, indicating less self-centered decision-making. Second, we show behavioral signatures that indicate specific disruptions of social basis function use in individuals with BPD. As in previous studies, we found that the primary basis function (reflecting the comparison own team vs other team) was balanced and symmetrical in NCCs (2,4). In BPD, on the contrary, it was significantly shifted: While individuals with BPD showed clear signatures of perceiving opponents as being grouped together in the same team (“them”), they did so significantly less for the grouping of themselves with their own partners (“us”), manifesting as outgroup overweighting. Using smartphone-based ecological momentary assessment, we found that social basis function use flexibly regulates momentary loneliness according to fluctuation in self-evaluation in daily life. This regulatory flexibility is disrupted in individuals with BPD. Together, these findings suggest that individuals with BPD have an unbalanced representation of social structure and a more isolated sense of self.

**Table 1:** Demographics of BPD individuals and non-clinical controls (NCC). Demographic and clinical characteristics of individuals with borderline personality disorder (BPD) and non-clinical control participants (NCC). Values are reported as mean (standard deviation, SD) unless otherwise indicated. Age range is given in years. Gender is reported as number of participants: W = women, M = men, TM = trans men. BPD criteria refer to the number of diagnostic criteria for borderline personality disorder met according to structured clinical assessment. BSCL Sum Score = Brief Symptom Checklist total score, reflecting overall psychopathological symptom severity. We included only NCCs without psychiatric diagnosis who were not undergoing psychiatric or psychotherapeutic treatment (based on self-report). Cognitive ability reflects performance on a standardized cognitive assessment, the Matrix Reasoning Task (32) (MARS; see Methods). p-values indicate group differences between BPD and NCC.

| Measure | BPD | NCC | Group difference |
| --- | --- | --- | --- |
| Age | 18.49 (SD 2.73) | 18.66 (SD 2.84) | p = 0.769 |
| Age range | 13–25 | 13–26 | — |
| Gender | 41F, 4TM, 1M | 50F, 1M | — |
| BPD criteria | 5.41 (SD 1.86) | — | — |
| BSCL Sum Score | 26.87 (SD 5.83) | 13.11 (SD 3.60) | p < .001 |
| Cognitive ability<br>(MARS) | 27.57 (SD 5.92) | 27.04 (SD 5.23) | p = 0.643 |

## Results

### Abstract multi-person decision-making task and variables of interest

In a short (∼3min) pre-experiment, participants completed 120 trials of a of a standard random dot motion task (e.g. (33)). On every trial of this task, participants saw a random dot motion cloud with an average motion direction either towards the left or the right side. Participants were required to press the rightwards button to indicate rightwards movement, or the leftwards button to indicate leftwards movement. Initially, they received positive feedback for correct/successful responses (a yellow “coin”) and negative feedback for erroneous responses (a red “X”; Fig.1A). This feedback was hidden during the second half of trials. The pre-experiment established the meaning of these “performance cues” (yellow “coin”, red “X”) as successes and failures, respectively, and familiarized participants with the player locations on screen. Note that the random dot motion task itself was not of interest for the actual purpose of our experiment. It was merely a vehicle to establish the concept of “performance cues” and participants were told that they would be shown samples of these cues relating to themselves and other players in the main experiment. The pre-experiment thus prepared participants for the main experiment where they were shown performance cues for all players (in the absence of any random dot motion task)

**Fig. 1.**
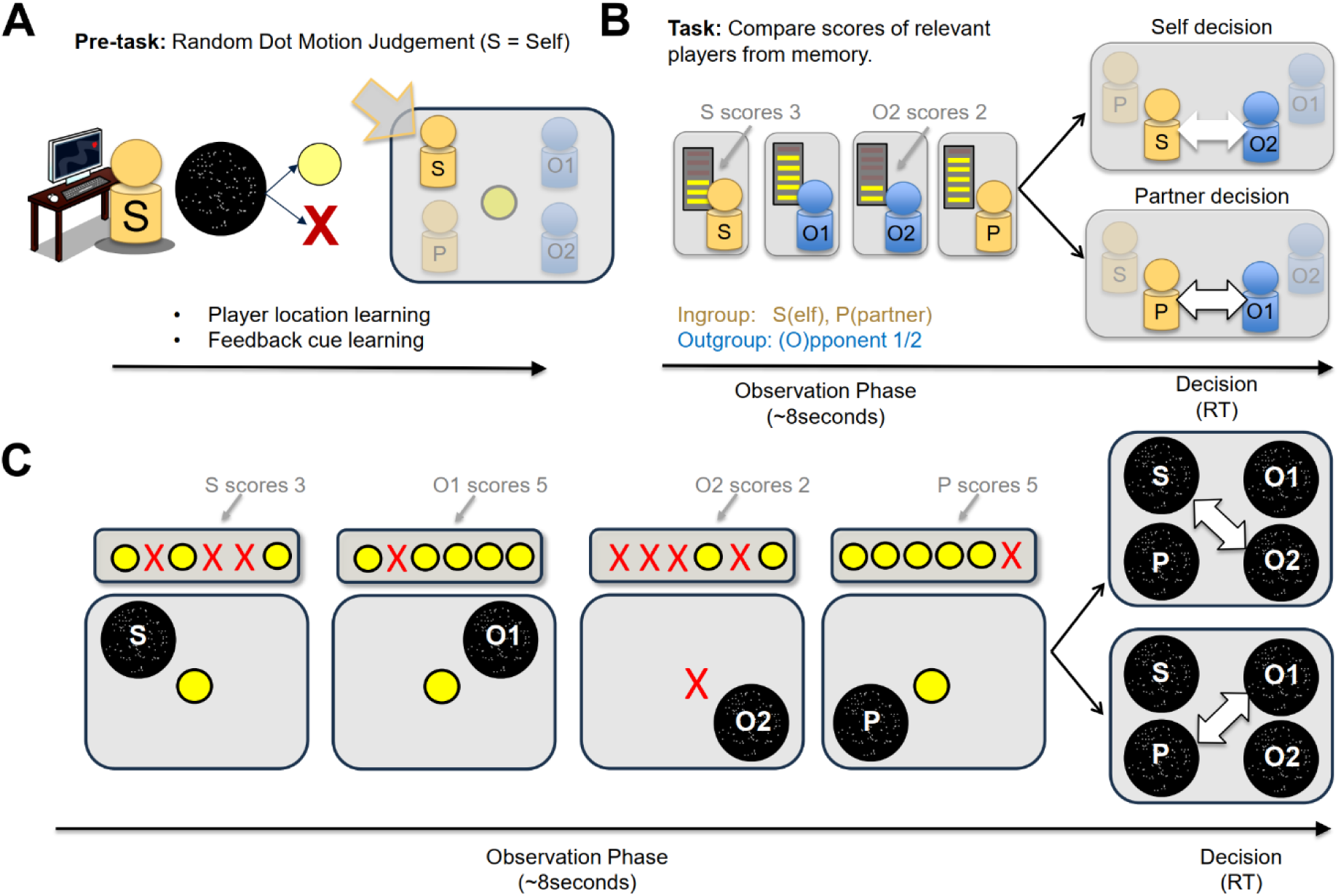
Abstract multi-person decision-making task. (**A**). **Pre-experiment.** As a pre-experimental task, participants performed a left/right motion judgment task with random dot stimuli, receiving feedback via yellow “coins” (correct) or red crosses (incorrect). Participants were paired with a partner (‘P’) against two opponents (‘O1’, ‘O2’) before starting the main experiment. **(B). Summary of trial timeline of main experiment.** The main task consisted of two phases per trial: an observation phase and a decision phase. During the observation phase, participants observed “performance cues” which looked similar to the pre-experiment feedback cues and related to the four players. Yellow coins indicated success and red crosses indicated failures and are symbolized here as horizontal bars in this panel. Participants needed to keep track of the overall number of success cues (i.e. the score) for each player (e.g., 3 for S, 2 for O2). In the subsequent decision phase, participants made one of two types of decisions indicated by an arrow cue: Self or partner decisions. The arrow cue indicated the relevant to-be-compared players. The two uncued players were decision irrelevant and needed to be ignored (greyed out players). In self-decisions, participants compared their own score (‘3’ in this example) with the score from the indicated opponent (‘2’ in this example, for O2). In partner decisions, they compared their partner’s score with the score of the indicated opponent. Note that both S and P were equally often paired with O1 and O2. They responded with a left/right button press congruent with the side of the alleged higher performer. **(C). Detailed trial timeline of main experiment.** The Panel illustrates some details of the summarized trial timeline shown in panel B. Note that the players were indicated as static random dot motion clouds to tie in with the pre-experiment. For the participant, initials were overlaid on the dot clouds. Scores of participants were presented as a series of 6 performance cues where the number of yellow coins indicated the player’s score. The performance cue order for each player was random, and only the summed scores mattered; different cue orders for the same score were irrelevant. At decision time, an arrow indicated which players were meant to be compared by pointing towards their respective dot clouds with initials.

In the main experiment, participants performed an abstract multi-person decision-making task (Fig.1B). At its core, the task was a social working memory task where participants on every trial remembered players’ performance scores. Critically, a social structure was established by grouping players into two teams, one comprising the participant (Self, S) and a partner (Partner, P) they were cooperating with. This team (i.e., S and P) is the “ingroup”. The “outgroup” comprised two other players that S and P were competing with (opponent 1, O1 and opponent 2, O2). On each trial, in an initial observation phase, a sequence of six performance cues linked to each player was shown (Fig.1B,C). The cues were visually identical to the cues used in a pre-experiment, such that participants were familiar with their meaning: Yellow ‘coins’ indicated “successes” and red crosses indicated “failures”. Participants’ task in this observation phase was to keep track on the score (i.e. the sum of successes, e.g., 4 out of 6) per player. Note that participants were not able to control which cues would be shown, they simply had to keep track of which cues were shown. In a subsequent decision phase, participants’ task was to make decisions resulting from a comparison of two of these scores from memory. An arrow cue indicated which players’ scores should be compared. These comparisons included either S and one of the Os (each with same probability) during ‘self decisions’ or P and one of the Os (each with same probability) during ‘partner decisions’. Participants were instructed to indicate the player with the higher score with a leftwards or rightwards button press in the direction of the chosen player. We denote the relevant, to-be compared opponent as Or (Opponent-relevant) and the irrelevant opponent as Oi (Opponent-irrelevant). Self and partner decisions occurred with the same frequency and pseudo-randomly.

The experimental paradigm therefore required participants to make decisions about information assigned to themselves (S) and specific players they held cooperative (P) and competitive relationships (O1, O2) with. Importantly, this allowed us to examine how participants represent social structure. It allowed us to extract factors of interest that were specifically social. The first is the factor “decision type”, namely whether decisions were made on behalf of oneself or the partner. Importantly, self and partner decisions were equally frequent and difficult, and correct decisions in either earned the participants points – which was the goal of the task. This meant that, from a normative neuroeconomic perspective, participants should perform equally well in self and partner decisions. Any differences between self and partner decision would indicate different processing of relational information that was tagged onto either oneself (S) or a cooperator one was acting on behalf of (P).

The second factor of interest is the factor “group”. Our experimental paradigm allowed us to compare how decisions were impacted both by members from the ingroup (S, P) and by members from the outgroup (O1, O2). Again, if done in an “optimal” way from a neuroeconomics perspective, information from ingroup and outgroup should be weighted identically. Any asymmetries in this weighting, however, would indicate socially specific effects that relate to the cooperative and competitive structure that people are embedded in.

### Lower decision accuracy in individuals with BPD

We compared accuracy and the logarithm of the reaction times (logRT) across cohorts (BPD vs. NCC). We found that controls were more accurate in their decisions than BPD (t_95_=3.034; p=0.003). That is, NCC were more often correct when comparing performance scores of relevant players from memory (Figure 2). There was no significant group difference in reaction times (t_95_=1.686; p=0.095).

**Fig. 2.**
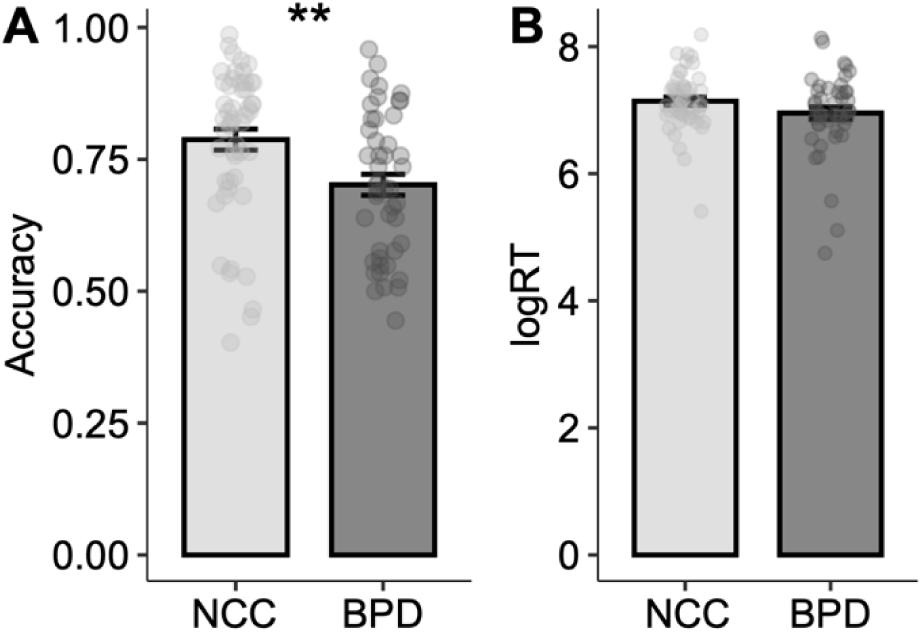
Individuals with BPD perform the task less accurately. (**A**). Accuracy (proportion correct decisions when comparing the performance of relevant players from memory) in BPD patients and controls. (**B**) No difference in logRT across groups. (N_BPD_=46, N_controls_=51). * p<0.05; **p<0.01; ***p<0.001; error bars are SEM.

### Reduced egocentric decision making in individuals with BPD

We went on to our key analyses of interest to determine how individuals with BPD’s decision-making are impacted by social structure. We specifically investigated how individuals with BPD weighted decision-related information in our task, focusing on the social dimensions of decision type (self vs partner decision) and group (ingroup vs outgroup). To do so, we conducted a regularized general linear model (GLM; see methods) analysis. We predicted choices in favor of the relevant player from the ingroup (S in self decisions, P in partner decisions), coded as 1/0. Importantly, we fitted the same regularization parameter for self and partner decisions to avoid bias. We regressed choices onto the performances scores of all four players. We coded the player identities as Self (S), partner (P), relevant opponent (Or; the opponent indicated by the arrow that should be considered in a decision), and irrelevant opponent (Oi; the member of the outgroup that is decision irrelevant). This model yielded “decision weights” reflecting the influence of each individual player’s performance score on participants’ choices. High decision weights indicate that a decision was strongly influenced by the very performance of a given player, whereas low decision weights indicate that the performance score of a given player did not influence participants’ decisions much. This allowed us to determine if participants weighed social information in a uniform way, or whether there were any socially specific distortions based on the factors manipulated by our experimental design, i.e. decision type (self/partner) or group (ingroup/outgroup).

First, we considered the decision weights of the relevant players that were to be compared on every dyadic decision (Fig.3A). This procedure resulted in comparisons of four decision weights, two for self decisions (S and Or), and two in partner decisions (P and Or; Fig.3A). We sign-adjusted the outgroup decision weights to allow numerical comparisons across ingroup and outgroup (by multiplication with “-1”; see Methods). Note that in case participants did not assign any social meaning to the different players, we would expect all decision weights to be the same and uniformly positive – for self and partner decisions and for ingroup and outgroup weights (Fig.3 B,C).

**Fig. 3:**
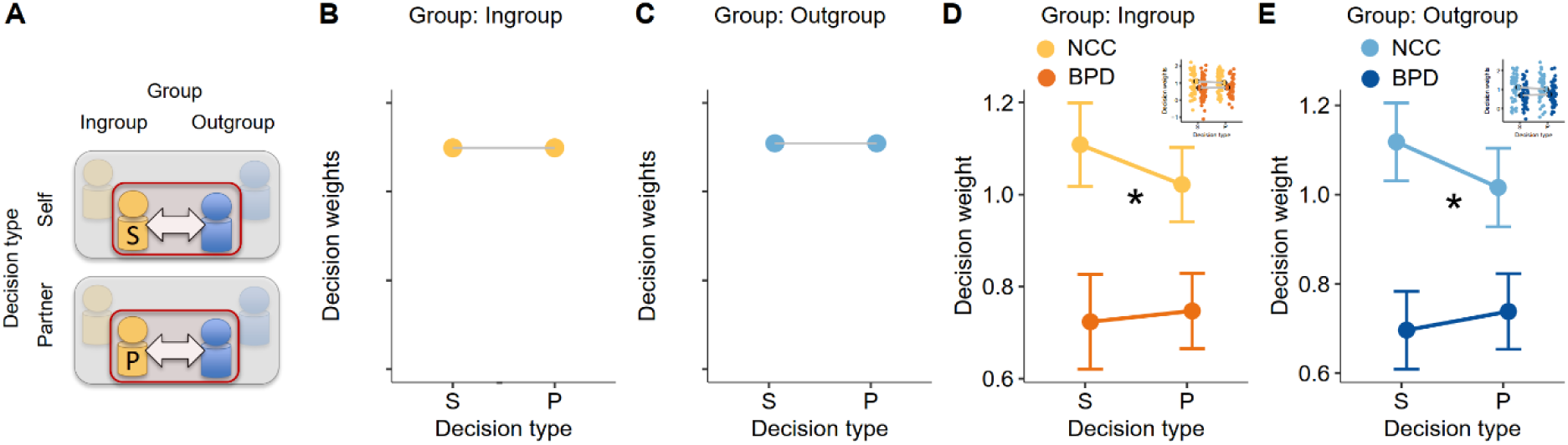
Less egocentric use of relevant decision information in individuals with BPD. (**A**). We compared the decision weights for relevant ingroup and outgroup members across self and partner decisions. Red boxes indicate decisions weights of analyzed players. Greyed out players are irrelevant for the decision at hand. (**B**). Schematic of decision weights in the absence of any social modulation: Ingroup members should have a positive weight on decisions and this should be equal for when S needs to be considered in a self decision and when P needs to be considered in a partner decision. (**C**). Analogously, the weights of the opponents should be identical to the ingroup weights, no matter if it is a self decision or a partner decisions. (**D,E**). Measured decision weights for the relevant player from one’s own group (panel D) and the relevant player, Or, from the other group (panel E). Our obtained results differed from this uniform pattern, and we found instead that decision weights differed between NCCs and individuals with BPD along the social dimension of group. While NCCs had reduced decision weights for relevant players in self compared to partner decisions, this was not the case for individuals with BPD (significant cohort x decision type interaction). The effect was stable over ingroup (panel D) and outgroup (panel E), i.e., there was no interaction with group. (*, p<0.05; significance star indicates interaction of decision type with cohort; N_BPD_=46, N_controls_=51)

We compared the decision weights of the relevant players using a 2 [decision type: self, partner decision] x 2 [group: ingroup, outgroup] x 2 [cohort: BPD, NCCs] mixed effects ANOVA. The analysis showed a main effect of cohort (F_1,95_=9.264; p=0.003) indicating higher decision weights for NCC compared to BPD patients. That is, across all relevant players their actual performance scores influenced decisions in NCC more than in BPD. As higher decision weights indicate higher precision in using decision-relevant information for choice, this finding is in line with the result of our descriptive analysis above showing higher decision accuracy in controls. However, importantly, we found a significant interaction of cohort with one of our social dimensions of interest, namely decision type (F_1,95_=4.474; p=0.037; Fig.3D,E). Figure 3 D-E reveals that while in NCCs decision weights were relatively higher in self-, compared to partner-decisions, BPDs decision weights were reversed and higher in partner compared to self decisions. Post-hoc analyses showed that this interaction effect of cohort x decision type was driven by a significant effect of decision type in NCCs (significant main effect of decision type in 2 [decision type] x 2 [group] ANOVA; F_1,50_=4.964; p=0.030) which was absent in BPD (main effect of decision type in 2 [decision type] x 2 [group] ANOVA; F_1,45_=0.595; p=0.444; Bayesian evidence for the null: BF_01_=4.141).

Our observation that in NCCs decision-weights increased when decisions are made about the Self in is in line with a vast literature in neurotypical participants that people have improved decision making about self-related compared to other-related information and this effect has been interpreted as a ‘self-bias’ or egocentric decision-making (4,34,35). The interaction effect of cohort with decision type may therefore be interpreted as a significantly reduced “self-bias” in individuals with BPD, reflected in a weaker preferential weighting of self-relative to partner-related information during decision-making. None of the other effects in the ANOVA were significant: notably, there was no significant interaction between cohort (NCC vs BPD) and group (F_1,95_=0.077; p=0.781). There was also no triple interaction between decision type x cohort x group (F_1,95_=0.082; p=0.775). The latter indicates that the cohort effect on decision type we report is not limited to the decision weights for ingroup members, S and P (Fig 3D), but also extends to the relevant outgroup members, the Os (Fig.3E): NCC showed a self-bias in the weighting of social information when making decisions on behalf of themselves compared to on behalf of their partner, and in doing so, they not only weighted information about themselves more strongly, but also information about the Or. In individuals with BPD by contrast, the self-bias was abolished across both in- and outgroup conditions. The latter finding is not compatible with a strict interpretation of a self-bias as exclusively relating to self-associated information in NCC, as it extended to Or. Instead, it suggests a more general advantage in information weighting in NCC that extends to various types of information as long as they are to be combined and weighted in relation to themselves. Individuals with BPD, on the other hand, did not show this egocentric bias.

### Disrupted balance of social basis functions in individuals with BPD

After showing altered patterns of social decision making in individuals with BPD when tagging information onto social identities such as oneself and the partner, we went on to examine potential disruption in the flexible representation of social structure more generally. To do so, we focused on the decision weights of the irrelevant players. We did so for the following reason. We have shown that the team structure embedded in our task - cooperating with a partner against two opponents – reflects a fundamental social structure that acts as a mental scaffold – a primary basis function (2,4). This grouping structure is reflected in subtle effects of irrelevant players on decisions in line with group membership: high scores of an irrelevant ingroup partner increase participants’ likelihood to choose their own team, while high scores of irrelevant outgroup opponents increase participants’ likelihood to choose the opponent team. Thus, although these players are not directly relevant for the current decision, their information is incorporated according to the underlying social structure, indicative of a pairwise mental grouping of S and P, and Or and Oi – the primary basis function. The weighting of irrelevant players therefore provides a window into the flexibility with which individuals adjust and integrate social representations beyond the immediate decision context. We therefore investigated strength and directionality of irrelevant player’s decision weights.

We analyzed the decision weights from the above regularized regression, focusing on the influence on irrelevant players’ performance on decisions (Fig.4A). Specifically, we analyzed the decision weights associated with the identities of P and Oi in self-decisions, and S and Oi in partner decisions. Following the same analysis rationale as above, we analyzed the irrelevant decision weights with respect to the social dimensions of decision type (self vs partner decision) and group (ingroup vs outgroup). As above, we sign-reversed Oi’s decision weights by multiplying them with “-1”. Positive decision weights of irrelevant partners and Ois were thereby comparable and indicate equally strong influences of these players in the direction of their respective team. Similar to the analysis above, we would expect uniformly positive decision weights that are equal for decision type and group if participants only considered the functional role of the presented performance information irrespective of their social identities (Fig.4B,C). However, again, we found that individuals with BPD and NCCs significantly differed regarding a social dimension of the task, as outlined in the next paragraph.

**Fig. 4:**
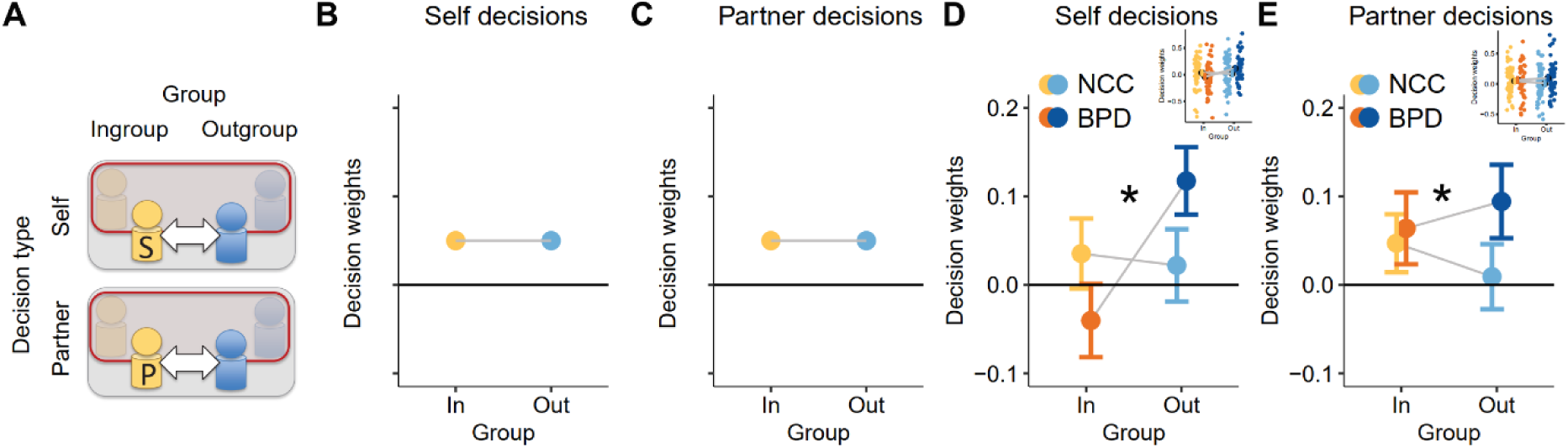
Individuals with BPD displayed shifted balance of irrelevant player’s decision weights. (**A**). We compared the decision weights for irrelevant ingroup and outgroup members across self and partner decisions. Red boxes indicate decisions weights of analyzed players. Greyed out players are irrelevant for the decision at hand. (**B**). Schematic of decision weights in the balanced social basis function use: Ingroup members should have a positive weight on decisions and this should be equal for irrelevant ingroup and outgroup members alike when decisions are made about oneself. (**C**) Analogously, both types of irrelevant weights in partner decisions should be identical to self decisions and should be the same for irrelevant ingroup and irrelevant outgroup members. (**D,E**). Our obtained results differed from this predicted pattern, and we found instead that decision weights differed between controls and BPD patients along the social dimension of group. BPD patients showed an increase in decision weights for outgroup members relative to ingroup members, while controls showed balanced effects across ingroup and outgroup members. This indicated a disruption in the use of the primary basis function in individuals with BPD. The effect was stable over self decisions (panel D) and the partner decisions (panel E). (*, p<0.05; significance star indicates interaction of group affiliation with cohort; N_BPD_=46, N_controls_=51).

We compared the decision weights of the irrelevant players using a 2 [decision type: self, partner] x 2 [group: ingroup, outgroup] x 2 [cohort: BPD, NCCs] mixed effects ANOVA. We found an interaction of cohort with group (F_1,95_=4.414; p=0.038; Fig.3D,E). That is, a balance of decision weights of irrelevant ingroup and outgroup members was systematically different in BPDs compared to NCCs. This interaction effect was explained by the fact that, in individuals with BPD, outgroup members being weighted more positively than ingroup members (main effect of group in 2 [decision type] x 2 [group] post-hoc ANOVA in BPD; F_1,45_=5.882; p=0.019). The above interaction effect indicated that this was significantly different in NCCs, who did not show a significant difference in decision-weights between irrelevant ingroup and outgroup members (absent main effect of group in 2 [decision type] x 2 [group] post-hoc ANOVA in NCC; F_1,50_=0.384; p=0.538; Bayesian evidence for the null: BF_01_=3.365). Again, this effect cannot be explained in terms of better overall decision performance of controls. In sum, this observed asymmetry in BPD is indicative of disrupted primary basis function use and a diminished tendency to embed the four players into a self-partner vs O1-O2 group structure. While BPD grouped up information when it concerned their opponents, they expressed a joined group representation less when they themselves were included in the team, a phenomenon we term outgroup overweighting. In essence, this can be interpreted as a more isolated sense of self in BPD. This aligns with clinical observations in BPD of chronic feelings of not belonging, even within close relationships where others might be expected to be represented as distinct and meaningful individuals.

### Outgroup overweighting predicts daily loneliness via momentary self-evaluation

We investigated real-world consequences of laboratory-assessed altered social basis functions. To this end, we focused specifically on “outgroup overweighting”, the asymmetry of decision weights between irrelevant ingroup and irrelevant outgroup members (sum of decision weights for irrelevant outgroup members minus sum of decision weights or irrelevant ingroup members). This variable captured precisely the imbalanced weighing of social structure in individuals with BPD shown in the previous section (Fig.4). Compared with NCCs, individuals with BPD exhibited greater “outgroup overweighting” (t_95_ = -2.101; p=0.038, Fig.5A). In line with our interpretation of this representing a more isolated self, we used this signature to predict daily loneliness reports we collected using smart-phone based ecological momentary assessment (EMA). Individuals with BPD are known to experience excessive chronic loneliness compared to both non-clinical controls and even those with other psychiatric disorders (13–16). Note that this analysis not just established links between experimental variables and one-shot self-assessments, but it allowed us to relate outgroup overweighting specifically to fluctuations in moment-to-moment self-assessments in participants’ daily life over a time period of 8 days.

**Fig. 5:**
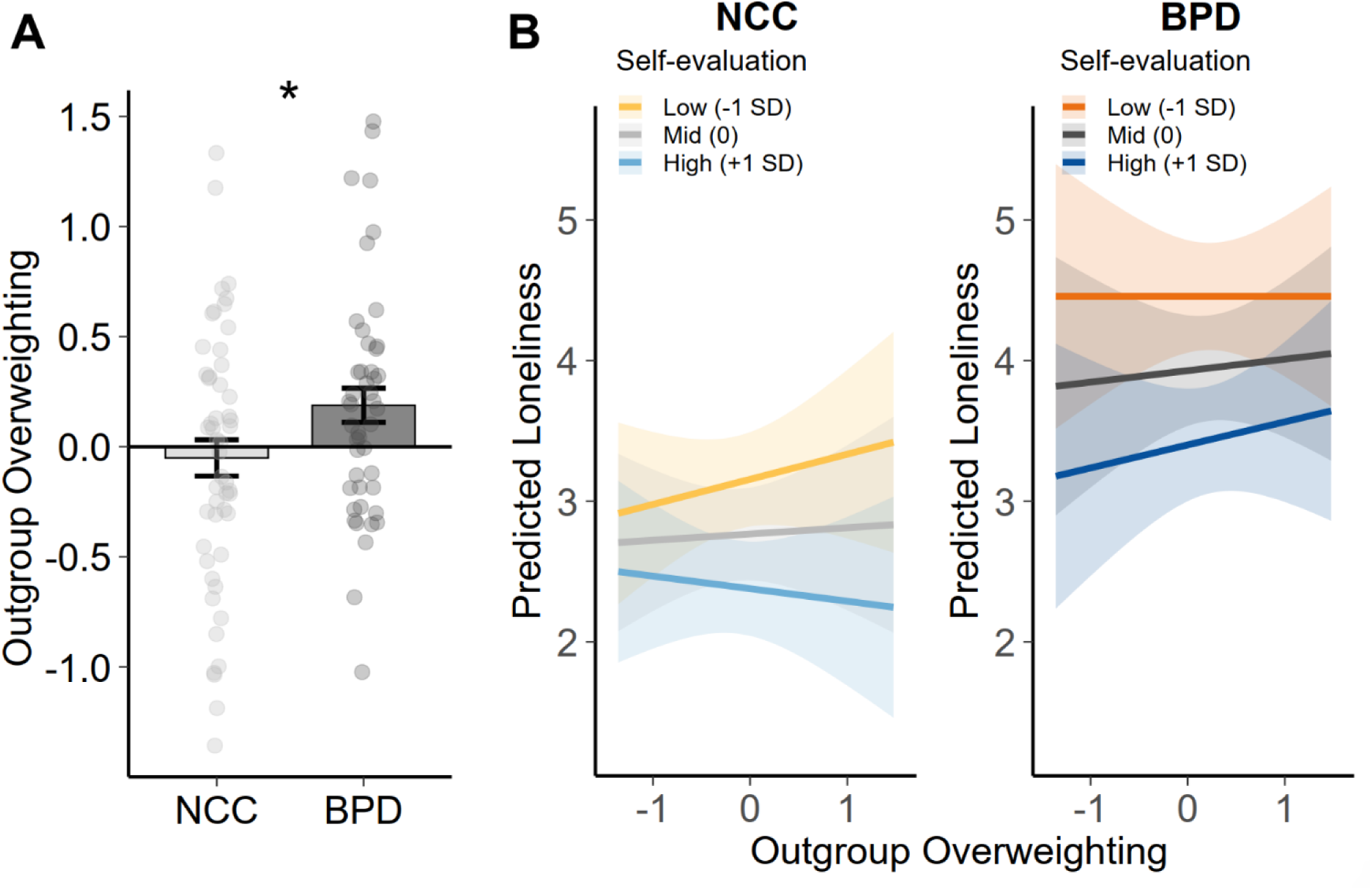
Individuals with BPD displayed shifted balance of irrelevant player’s decision weights. **(A)** The outgroup overweighting was compared between non-clinical controls (NCCs) and individuals with BPD. Individuals with BPD show significantly greater outgroup overweighting than NCCs, indicating a distorted social basis function representation, * p<0.05, error bars are SEM. **(B)** Regression slopes illustrating how outgroup overweighting predicted loneliness as a function of momentary self-evaluation. Individuals with BPD experienced greater loneliness than NCCs. In addition, we found outgroup overweighting showed a dynamic relationship with loneliness as a function of momentary self-evaluation in NCCs: greater outgroup overweighting was associated with higher loneliness when self-evaluation was low, but with lower loneliness when self-evaluation was high. The dynamic recalibration between outgroup overweighting and loneliness via self-evaluation was absent in individuals with BPD.

We employed a multilevel modelling approach linking between-person experimental data (outgroup overweighting) to within-person fluctuations in self-evaluation and loneliness measured via EMA (for details, see Methods). Participants’ momentary self-evaluation was assessed using the question “How good do you feel about yourself at this moment?”, while momentary loneliness was assessed using the statement “I feel lonely at the moment”, rated on a 1-7 Likert scale, during the same prompt. Outgroup overweighting, derived from the social group decision-making task, served as a stable between-person predictor as it reflected an experimentally measured individual difference in social basis function use. The model further included momentary self-evaluation, measured concurrently with loneliness via EMA, as a moderator. This was theoretically motivated by the architecture of social basis functions. Because social representations are referenced to one’s own position in the social world (4), self-evaluation may be seen as a self-referential anchor point through which outgroup overweighting is translated into subjective social experience. Momentary self-evaluation was person-mean-centered prior to analysis, yielding a within-person deviation score that captured momentary fluctuations in self-evaluation independent of stable interindividual differences in self-evaluation level. This centering approach ensured that a potential moderating role of self-evaluation would reflect genuine within-person dynamics rather than confounded between-person or group-level differences (which is particularly important given that BPD and healthy controls differ substantially in their average self-evaluation and loneliness levels) (36).

We tested a three-way interaction between outgroup overweighting, momentary self-evaluation, and cohort (BPD vs. NCC) predicting momentary loneliness, with model fit evaluated against a reduced model without the self-evaluation terms. To rule out a generic negative-affect account, we tested the specificity of self-evaluation as a moderator over and above momentary mood, which was also assessed via EMA. We observed a main effect of cohort (β = 1.174, SE = 0.310, t(91.80) = 3.78, p < .001), namely that BPD patients reported elevated levels of loneliness, a pattern in line with many previously published results (13–16). The model also revealed that momentary self-evaluation was associated with loneliness ratings (β = −0.389, SE = 0.031, t(2035.0) = −12.49, p < .001) and this was moderated by cohort (β = −0.139, SE = 0.048, t(2045.72) = −2.92, p = .004). Self-evaluation showed a stronger association with loneliness in BPD (β = −0.529, SE = 0.036, p < .001) than NCC (β = −0.389, SE = 0.031, p < .001), and these slopes differed significantly between cohorts (contrast: b = 0.139, SE = 0.048, t(2051) = 2.91, p = .004). This pattern is consistent with the notion that in BPD, where the social basis function fails to embed the self within a coherent team structure, momentary self-evaluation becomes the primary available signal for felt social belonging, rendering loneliness tightly coupled to transient fluctuations in self-perception.

Further, we found a two-way interaction of momentary self-evaluation and outgroup overweighting on loneliness (β = −0.134, SE = 0.051, t(2040.39) = −2.60, p = .009). Most interestingly, the mixed model revealed a three-way interaction between outgroup overweighting, self-evaluation, and cohort (β = 0.217, SE = 0.082, t(2020.53) = 2.63, p = .009). In a post-hoc test, we re-ran the model separately for both cohorts. This showed that the interaction between self-evaluation and outgroup overweighting was significant in NCC (β = - 0.135, SE = 0.045, t(1126.77) = -2.974, p = .003), but not in BPD (β = 0.076, SE = 0.072, t(757.80) = 1.057, p = .291). Thus, in NCC the association between outgroup overweighting and loneliness reversed in sign across levels of momentary self-evaluation: it tended to be positive during moments of low self-evaluation and negative during moments of high self-evaluation, consistent with the significant interaction (Fig.5B). This suggests that outgroup overweighting, which reflects a disrupted primary basis function in which the self fails to find its appropriate team-embedding, is not inherently pathological. Rather, its translation into subjectively experienced loneliness depends on the momentary self-referential context.

## Discussion

BPD is characterized by profound disturbances in self- and other-related functioning and in the formation of interpersonal relationships. We have recently shown that people represent social basis functions – basic patterns of social relationships such as the difference structure between self and others (2–4). Here, we used an abstract multi-person decision-making paradigm to investigate how young individuals with and without BPD flexibly represent social structures. Specifically, we examined whether social structure representations are disrupted in BPD compared to NCC, and whether they relate to real-world loss of social structure – such as loneliness assessed using EMA. We report three key results: First, individuals with BPD did not show an egocentric decision-making bias typically observed in NCCs, with no improvement in decision weighting for self-compared to partner-related information. Second, participants with BPD showed an imbalanced weighting of social structure, reminiscent of outgroup overweighting: they displayed a pronounced tendency to group opponents together, yet they were less inclined to form groups that included themselves. Third, we analyzed these indices of disrupted social basis functioning in relation to repeated smart-phone based ecological momentary assessment of loneliness and self-evaluation in daily life. We found that outgroup overweighting predicted momentary loneliness in daily life, and exclusively so when moderated by momentary self-evaluation in NCC. However, this flexibility modulation between self-anchor and outgroup-overweighting in relation to loneliness was absent in individuals with BPD.

### Reduced egocentric decision-making in BPD

Our observation that decision weights increased for self-compared to partner-related decisions is consistent with a well-established self-bias in human cognition, whereby self-related information is processed with preferential precision across multiple domains including (working) memory (37,38), perception (39), learning (34,40), and decision-making (4,35). Once learned, self-related beliefs tend to persist despite disconfirming evidence (41). Indeed, such self-reference was suggested to be adaptive, in its role as an integrative mechanism, or “associative glue” facilitating the binding of heterogeneous information types across multiple stages of cognitive processing (42). This may enable us to build more accurate models of the self and those similar to us (41,43).

In line with this interpretation is the observation in our study that decision weights were increased in self-decisions for both self and the relevant opponent. This finding is difficult to reconcile with a strict interpretation of self-biases as operating exclusively on self-associated information. It is, however, consistent with the idea that self-referencing acts as an “associative glue” that increases precision for self-decision-relevant information more generally. Notably, disturbances in the sense of self, including chronic instability in self-image, values, and goals , are among the core diagnostic criteria of BPD (44,45), making self-referential processing a particularly relevant target of investigation in this population (8,46–48). Consistent with this, participants with BPD showed a significant reduction in the self-relevance advantage alongside generally lower decision accuracy. This finding aligns with, and may help explain, the well-documented disturbances in self- and other-representations characteristic of BPD (5,23,24,49,50). If self- and other-representations are unstable or insufficiently differentiated, the precision advantage for self-related information observed in neurotypical controls may be lost in BPD. This is consistent with broader clinical accounts of the disorder as marked by blurred self-other boundaries and chronic instability in self-representation.

### Outgroup-Overweighting as a cognitive signature of an isolated Self in BPD

Our multi-person experimental approach allowed us to further examine how representations of self and others were embedded into the social structure of two competing teams (self and partner versus two opponents, i.e. “us vs. them”). In our past work, we found that people use a so-called “primary basis function” – the difference between own and opponent team – as a mental scaffold that enables both efficient and flexible decision making (2–4). This aligns well with ingroup and outgroup being salient social categories (51–54), but reinterprets these accounts from an efficient coding perspective (27). Here we replicate these findings in NCC. In NCC, irrelevant teammates and irrelevant opponents exerted balanced and opposing influences on decisions, consistent with an intact mental grouping of the four players into “us vs. them”, the primary social basis function. This structure organizes social information around the self as an anchor point, such that the partner is implicitly recruited as an ally and opponents as a contrasting outgroup. In individuals with BPD, this balance was disrupted which manifested as a shift in irrelevant players’ decision weights: Individuals with BPD showed reduced weights for their own team and elevated weights for the opposing team. This indicates that players from one’s own team were less “grouped together”, while players from the other team were grouped together even more. Irrelevant opponents exerted disproportionately stronger influences than irrelevant partners, indicating that the self fails to recruit its partner into a coherent team representation. We refer to this as outgroup overweighting, a signature of a disrupted primary basis function in which the self lacks a stable social anchor. Indeed, our previous work showed that social basis function develops across adolescence (2), a developmental period that temporally overlaps with increased vulnerability to psychiatric conditions, including the onset of BPD (55,56). This developmental overlap raises the possibility that altered social basis function could be explored as a potential behavioral signature associated with BPD vulnerability.

Importantly, the increase in outgroup overweighting in people with BPD demonstrates that basis function use can be fundamentally socially configured by the place of oneself within the pattern of social relationships. The direction of basis function points towards a more isolated self in people with BPD, where alliances are perceived strongly among members of other social circles (“them”), but not towards one’s own ingroup (“us”). This helps explain common findings in experimental and observational studies of BPD patients feeling excluded (for a meta-analysis see (57)). Notably, pronounced feelings of exclusion are evident in BPD even in scenarios where participants are explicitly included (57–59). Previous accounts have explained this by increased rejection sensitivity (58,60,61) or negatively distorted interpretations of neutral or even positive social experiences (20,62,63). Our study offers a potential cognitive mechanism: disrupted social basis functions that prevent individuals with BPD from forming a stable mental representation of themselves as embedded within a social group, leaving the self without a team-anchor, and rendering social belonging chronically uncertain. Indeed, our findings might also help explain the mechanics underlying modern psychotherapies for BPD. In mentalization-based treatment (64), re-establishing a so-called “we-mode” in BPD patients is a key goal (65). In dialectical behavioral therapy, dialectical strategies aim to foster a collaborative bond between therapist and client (66).

### Associations of basis functions with momentary-assessed loneliness

To connect these mechanistic findings of self-other interactions to subjective experience, we leveraged repeated ecological momentary assessment, examining how individual differences in outgroup overweighting relate to momentary loneliness and self-evaluation in daily life. Note that although NCCs showed no systematic outgroup overweighting at the group level, there were individual differences in this index also in NCCs. We observed a crossover interaction in healthy controls: outgroup overweighting predicted a tendency towards greater loneliness during moments of low self-evaluation. This observation refines neuro-cognitive theories of loneliness, where a perceived gap between the self and others is defined as a central feature of loneliness (67–69). However, the same disposition was associated with reduced loneliness when momentary self-evaluation was high, a pattern that speaks to the context-dependence of social basis function disruptions. Outgroup overweighting may therefore not be inherently pathological, but instead reflect a tendency, the impact of which on subjective experience depends on a person’s momentary sense of self. For example, imagine someone at a party who tends to perceive other social circles as more tightly knit than their own: when feeling confident, this observation may simply feel like social clarity - “that’s their group, I know where I stand.” When feeling insecure, the very same perception may become “I don’t belong here, I feel lonely”.

In BPD, this dynamic flexibility was absent, an observation consistent with findings of chronically elevated levels of loneliness and social isolation in this patient group (13,70–72), as well as stickiness in updating social priors (20). This interpretation resonates with modern accounts of loneliness that emphasize the role of distorted internal models of the self and others in perpetuating subjective feelings of loneliness (73): Outgroup overweighting may be understood as a negatively skewed prior about one’s position in the social world that biases the interpretation of social experiences toward exclusion. In BPD, where the prior is more severely distorted and self-evaluation chronically low, this mechanism may contribute to the self-perpetuating nature of loneliness in the disorder.

### Conclusions

Our findings suggest that individuals with BPD differ not only in how they represent themselves and others as individuals, but also in how they represent the broader social structures in which these relationships are embedded. Using an abstract multi-person decision-making paradigm, we identified disruptions in social basis function use characterized by reduced egocentric decision-making and outgroup overweighting, indicating a diminished tendency to mentally embed the self within a cooperative social group. In other words: a pattern characterized by an attenuated sense of “we” alongside a heightened salience of “them.” Importantly, these experimentally measured alterations in social structure representation were linked to momentary experiences of loneliness in daily life through ecological momentary assessment. Together, these results provide a mechanistic account of disturbed self–other functioning in BPD and suggest that altered representations of social structure may contribute to feelings of social disconnection and isolation.

## Methods

### Sample Description and Recruitment

Our sample comprised 104 participants. We excluded 7 participants because their logfiles were corrupted or incomplete. The final sample comprised 97 participants (51 non-clinical controls [NCCs], 46 individuals with BPD). The Ethics Committee of the University Hospital Würzburg approved the study (#29/21-AM, accepted 05.03.2021). All participants (and caregivers for those under 18) provided written informed consent prior to study participation. Fluency in German was required for inclusion. All participants took part in a larger study that included a virtual reality paradigm (74), a battery of cognitive experiments, Ecological momentary assessment, and questionnaires, the results of which have been, or will be, reported separately (20,74). Participants were compensated with 120 € for study participation. Before completing the task reported here, participants were informed that they could gain a bonus depending on task performance. This additional bonus ranged between 1 and 2 €. Note that in order to accommodate patients’ preferences and to reduce their burden associated with study participation, some participants completed the study online, and others did so in-person. Importantly, setting (online vs. in-person) was matched on an individual level, that is, if an individual with BPD completed the task in person, the matched NCC did so as well and alias for the online setting. In that way we ensured that group differences cannot be due to different settings Further, even the task instructions were fully computerized, and instructions were delivered via the same computer on which participants performed the task. Therefore, the experience of doing the experiment itself, including the social framing, was identical for in-person and online data acquisition. Individuals with BPD and NCCs were recruited as follows.

### Individuals with BPD

Adolescents and Adults aged 13 to 25 years with symptoms of BPD were recruited from the clinical population. Consistent with the typical gender distribution observed in clinical populations in Germany (e.g., (75) and population-based studies of BPD (76,77), the majority of participants recruited for this study were female (44 out of 45 were born as women; see Table 1). All participants were either currently in treatment or had received treatment for BPD previously. Treatment modalities included inpatient and outpatient psychiatric care, psychotherapy, or specialized therapeutic residential programs for BPD, based on a dialectic behavioral therapy approach (DBT-A) (78). Collaborating centers included the Departments of Child and Adolescent Psychiatry and Adult Psychiatry at the University Hospital Würzburg, inpatient DBT centers in Lübeck and Marsberg, and therapeutic DBT residential programs in Germany. To be included in the study, participants underwent a standardized diagnostic assessment (SCID-II), conducted by a study psychologist, confirming that patients were not fully remitted, but that at least three BPD criteria were met at the time of participation. In adolescence, 3 BPD criteria is considered as clinically meaningful (79,80). This diagnostic procedure was performed following informed consent. Participants who did not meet the minimum threshold of three criteria were excluded. As is typical in BPD, most participants self-reported comorbidities. The most frequently reported comorbidities were depression (n=43), eating disorders (n=12), and post-traumatic stress disorder (n=11). Regarding psychotropic medication, participants reported the following: no medication (n=20), one medication (n=15), multiple medications (n=18). Medication types included SSRIs and SNRIs (n=24), neuroleptics (n=16), stimulants (n=6), sleep medication (n=8), and TCAs (n=5).

### Non-clinical control participants

Non-clinical Control participants (NCCs) were recruited from the participant database at the University Hospital Würzburg and via flyers to individually match BPD participants in age, sex and data acquisition setting (online vs. in-person). Mental and neurodevelopmental disorders were excluded based on self-report prior to inclusion and reconfirmed through the questionnaire battery. Included NCCs reported no current or past psychiatric diagnosis and no history of psychotherapeutic or psychiatric treatment. None of the NCCs were taking psychotropic medication. Consistent with the recruitment procedure, the Brief Symptom Checklist (81) sum score (indicating general psychopathology symptoms) was substantially lower in the NCC cohort compared to the BPD cohort (see Table 1).

### Experimental procedures

At the beginning of each testing session, participants performed a computerized task that comprised a pre-experiment and the main group decision-making task. The task (including pre-experiment and main decision-making experiment) was programmed in jspsych (82).

#### Pre-experiment

Participants performed a pre-experiment involving random dot kinematogram (RDK) stimuli. Participants judged the motion direction of RDK stimuli, presented using the Variable Coherence Random Dot Motion (VCRDM) toolbox (version 2; https://shadlenlab.columbia.edu/resources/VCRDM.html). The participants pressed left/right buttons to indicate congruent leftwards/rightwards motion directions of the RDK stimuli. Participants were made aware that they would perform these motion judgments at varying levels of coherence making the detection of the RDK stimuli easier or more difficult. The cues used to indicate correct and incorrect RDK performance (a yellow ‘coin’ and a red ‘X’) were the same ones used to indicate correct and incorrect performance in the subsequent main experiment, the group decision-making task. After the pre-experiment, participants went on to the main group decision making experiment.

The pre-experiment comprised 120 RDK trials using motion coherences, with 0.512 for 20% of the trials and 0.032 for 80% of the trials. A coherence of 0.512 represented an easier motion direction judgment. Each RDK stimulus was presented for a maximum duration of 1 sec, requiring participants to make their decisions within that time frame. Failure to respond within this time window was counted as incorrect performance and indicated by a “missed!” message on the screen. Participants were made aware of this and instructed to avoid ‘missed’ trials. The pre-experiment had two sub-parts: one with feedback and the other without. The pre-experiment took approximately 3 minutes.

#### Group decision-making task

Right after the pre-experiment, participants performed the main group decision making experiment for approximately 30 minutes. That is, they played a computerized task in which they observed the same type of performance cues that they had been familiarized with in the pre-experiment (yellow ‘coins’ and red ‘X’s). They were told that they would be observing performance cues linked to the four different players’ performance in the pre-experiment, themselves and three other players (4). The players were divided into two teams: the participant (“S”) and their partner (“Pa”) formed one team, and the other two players formed the opponent’s team (“O1”, “O2”).

The experiment comprised 72 trials. Each trial comprised an observation phase and a decision phase. During the observation phase, participants observed six brief performance cues per player, all presented centrally on the screen. These performance cues were identical to those used during the pre-experiment, with yellow ‘coin’ indicating successful performances and red ‘X’ cues indicating erroneous performances. At the same time, we used moving RDK indicating the relevant player related to the performance cue. For each player, each performance cue was presented for 300 msec with a 100 msec delay between them. During this time the player’s RDK was active to indicate the relevant player and the RDK movement ended precisely at the time the sequence of six performance cues also ended. To determine each player’s score, participants needed to keep track of the series of six performance cues, accumulating the yellow ‘success’ cues while disregarding the red ‘error’ cues. The number of successful performances, which ranged between 0 and 6, reflected the resulting scores. The performance cues for all four players were displayed in a random but counterbalanced sequence.

On each trial, after the observation phase, the decision phase followed. During the decision phase, participants retrospectively compared performance scores between players from their own team and the opposing team, using an arrow cue to indicate which players’ scores to compare. The decision was to choose the team that performed better, with each decision phase comprising two decisions based on the same set of performances in the trial. Each decision lasted until a response was given. Afterwards, a box appeared around the team that the participant had picked for 0.5 seconds. On each trial, two decisions occurred after each other with the restriction that it could not be exactly the same decision twice (e.g. not S vs O1). In contrast to previous versions of this experiment, no group decision existed and no “bonus” was utilized (see for context (2–4))

Two decision types were employed in the decision phase: self decisions and partner decisions. Both self and partner decisions occurred in equal number and pseudorandomly. In self decisions, the participant’s performance was compared with one of the two opponents. The performance of the other two players had to be ignored. In partner decisions, the partner’s performance was compared with the performance of one of the two opponents, ignoring the performances of self and the other, irrelevant opponent.

Participants’ incentive to do the task was to collect ‘points’ by making good decisions. If participants chose the player from their own team, then the points gained or lost were equal to the performance difference between the two cued players. If the one from their own team was indeed better, they received points equivalent to the true performance difference. Otherwise, they lost the points equivalent to the true performance difference. The outcome of choosing the player from the opponent team always led to a payoff of zero. This meant that participants were incentivized to choose the relevant player of their own team if that player had indeed performed better (in order to win points), and they were incentivized to choose the opposing player if the opposing player had performed better (to avoid losing points).

#### Ridge Regression analysis

We analyzed the data using MATLAB (83) and JASP (84). We fitted logistic general linear models (GLMs) to the choice data. All regressors were normalized (mean of 0, standard deviation of 1) and predicted the choice to pick the relevant player from their own team (=1). We performed regressions analyses for self decisions and for partner decisions. In both cases, the set of regressors comprised: the performance scores (i.e., numbers between 0 and 6) for the Self (S), the partner (P), the relevant opponent (Or), and the irrelevant opponent (Oi). We used ridge regression (85,86) to estimate the regressors’ decision weights (i.e., decision weights). Ridge regression penalizes large beta weights according to a regularization coefficient *λ* and thus prevents overfitting and improves generalization. We applied the regression model to all sessions using MATLAB’s lassoglm (setting Alpha to a very small value). First, we determined an appropriate regularization coefficient *λ*. To do so, we repeatedly fitted the GLM to each individual data set while varying *λ* between zero to 10^-3^ to 10^-1^ (log-spaced). During each fit, we used a three-fold cross-validation approach to determine the overall model deviance for each *λ* for all data sets combined. Herein, we also combined self and partner decisions to ensure that the resulting *λ* is equally determined by both trial types. We repeated this procedure twice. Finally, we selected the *λ* that resulted in the lowest overall model deviance over all participants and both trial types. This is the *λ* with the best cross-validated model fit, which was then used to run the ridge GLM of interest, for both self decisions and partner decisions. Importantly, the same best-fitting *λ* was used for all participants (patients and controls) and both conditions of interest (self and partner decisions) to enable fair statistical comparisons of beta weights within and across conditions.

Note that we sign-adjusted the resulting decision weights of the opponent players (Or and Oi) by multiplying them with “-1”. This enabled a clear comparison between ingroup and outgroup decision weights. Transformed in this way, if the decision weight of Or was, say, “2”, and the decision weight for the relevant ingroup member was also “2”, then this meant that the participant’s choice was equally driven by both types of information, albeit in the direction of choosing the other group in the former case and in the direction of choosing one’s own team in the latter case.

### EMA assessment

We conducted an 8-day Ecological Momentary Assessment (EMA) using the MovisensXR app on participants’ smartphones. To accommodate a VR experiment (74), for which EMA assessments needed to be completed on several days before and after participation, the assessment period was divided into a three-day and a five-day phase. As the MovisensXR app is only available for Android phones, we provided study phones for participants with iPhones. EMA involved participants responding to prompts and questions throughout the day using the app. Certain questions were asked 10 times a day, with intervals of 1.5 hours between 8 AM and 9:30 PM. Some questions were asked only 4 times a day at 9:30AM, 3:30 PM, 6:30 PM, 9:30 PM. Participants could reply to the prompt in a time window of 30 minutes. Among the broader set of questions asked (which will be reported elsewhere), here we focus on two probes, (1) participants’ momentary self-evaluation ratings and (2) ratings of loneliness. Self-evaluation was measured by the question “How good do you feel about yourself at this moment?”, asked 10-times a day and rated via a slider. To capture loneliness, participants were asked to rate the statement “I feel lonely at the moment” on a Likert scale from 1-7, with responses collected 4-times a day. As our analyses focused on the association between loneliness and self-esteem, only the prompts administered 4-times per day were included. Before data analysis, missing prompts, resulting from factors such as smartphones being switched off or compatibility issues with MovisensXR, were manually coded as missing value to allow for the calculation of the completion rate. No participants were excluded based on their completion rate, as our research question did not depend on the temporal structure of the data. The mean completion rate was 64.82% (SD = 20.59), with an average of 23.57 completed prompts per participant (SD = 8.24). Prior to modelling, momentary self-evaluation, mood, and tension were standardized within persons (i.e., centered on each participant’s mean and divided by their within-person standard deviation) to isolate momentary fluctuations from stable between-person differences. This standardization is undefined for participants who show no within-person variance on a given variable (a standard deviation of zero, i.e., an identical response at every prompt). Because the same analytic sample was retained across all models to permit valid model comparisons, observations were excluded if the within-person standardized value was undefined for any of the EMA state variables included in our models. This criterion excluded three participants who reported invariant momentary tension across the assessment period. The final analytic sample comprised N = 91 participants.

### EMA analysis: Mixed Effects Model

To examine whether outgroup overweighting predicts momentary loneliness in daily life, and whether this relationship is moderated by momentary self-evaluation, we employed a series of linear mixed-effects models (LMMs) using the *lme4* and *lmerTest* packages in R (87,88). All models were estimated using maximum likelihood (ML) to enable likelihood-ratio-based model comparisons. EMA and task data were available of 91 individuals in total, n=48 NCC and n=43 BPD patients (range of available ratings per person: 8-57).

The outcome variable was momentary loneliness as assessed via EMA (“I feel lonely at the moment.”). Outgroup overweighting, derived from the social group decision-making task, entered as a stable between-person predictor. Momentary self-evaluation, assessed via EMA at the same time point (“How good do you feel about yourself at this moment”), was within-person standardized prior to analysis (person-mean centered and divided by the within-person standard deviation), yielding a deviation score that reflects momentary fluctuations relative to each participant’s own baseline. This approach ensures that the moderating role of self-evaluation reflects genuine within-person dynamics, independent of stable between-person differences in self-evaluation level. The stable between-person component of self-evaluation (person mean, grand-mean standardized) was included as a covariate to partial out trait-level variance. All EMA variables were analogously within-person standardized. The day within the EMA period was included as a random slope to account for individual differences in loneliness trajectories over time.

Our primary model thus took the form:

*Loneliness ∼ Outgroup_overweighting × Cohort × Self-evaluation_within + Self-evaluation_between + (1 + Day | Participant)*

where *Cohort* (BPD vs. NCC) was entered as a between-person factor, *Self-evaluation_within* reflects momentary within-person deviations, and *Self-evaluation_between* is the stable trait covariate. The random effects structure included random intercepts and a random slope for day, without estimating their correlation to avoid overparameterization. Post-hoc analyses were conducted by re-estimating the primary model separately for BPD and NCC. Simple slopes were tested using the *emmeans* package (89). For additional control and sensitivity analyses, see Supplement.

## Acknowledgements

The authors wish to thank all participants for the time they invested without which this research would not be possible. We are grateful to all collaborating centers (LWL-Klinik für Kinder-und Jugendpsychiatrie Marsberg, JuLe Lübeck, Franz von Assisi gGmbH) for enabling recruitment and on-site data collection. We thank all student assistants for their help with data acquisition. The study was supported by grants from the German Research Foundation (DFG) awarded to AMFR (SFB 940/3 B7, RTG 2660-B2/project number: 433490190), as well as by funding from the Kaufmännische Krankenkasse (KKH, contact address: Kaufmännische Krankenkasse – Referat Prävention und Selbsthilfe, Mr. Tobias Bansen, Karl-Wiechert-Allee 61, 30625 Hannover, Germany). AMFR further acknowledges support from the German Research Foundation (DFG RE 4449/1-1), from grants by the Federal Ministry of Research, Technology and Space (BMFTR (CompExIn, FK 01GQ2302A, GAMKI-Wer wie was?, FK 16SV9364), and from a 2020 BBRF NARSAD Young Investigator Grant from the Brain & Behavior Research Foundation. MKW was funded by the UCL Institute of Mental Health, an UK Research and Innovation guarantee grant (UKRI; under the UK government’s HorizonEurope funding guarantee UKRI336 [selected as ERC Starting Grant, EC reference 1011159]) and a Medical Research Council grant (MRC; MR/Y010477/1). YL, SC report no financial disclosures.

## Author contribution

MKW.: Conceptualization, Methodology, Formal Analysis, Software, Writing – Original Draft, Supervision, Funding Acquisition. YL: Methodology, Formal Analysis, Software, Writing – Original Draft, Data curation. SC: Methodology, Formal Analysis, Software, Writing – Review & Editing. SM: Investigation, data curation, Writing – Review & Editing.MR: Investigation, , Writing – Review & Editing, Funding Acquisition. ASD: Conceptualization ,Writing – Review & Editing, Funding Acquisition. AB: Investigation, Writing – Review & Editing, Funding Acquisition, Project Administration. KG: Investigation, data curation, Writing, Formal Analysis, Software – Review & Editing. AMFR: Conceptualization, Methodology, Writing, Formal Analysis, Software – Original Draft, Supervision, Funding Acquisition, Project Administration.

## Code and data availability statement

The data and code supporting the findings of this study will be made publicly available upon publication.

## Supplement

### EMA: Control analyses

To assess the specificity of self-evaluation as a moderator, we additionally estimated a robustness model in which momentary mood and tension, also assessed via EMA, and within-person standardized, were included as fixed-effect covariates. This did not change the significance of our results of interest (significant three-way interaction cohort x momentary self evaluation on momentary loneliness p=.0099). We further tested a model in which mood replaced self-evaluation as the three-way moderator, using AIC to compare model fit. This indeed fit the data worse than the original model (ΔAIC = +29). To rule out that the between-person component of self-evaluation drives the moderation effect, we tested a model in which the person mean of self-evaluation entered as a full three-way interaction rather than as a covariate; however this more complex model did not improve fit (Δχ²(3) = 1.62, p = .655, ΔAIC = +4.4), confirming that the moderation is specific to within-person fluctuations.

## References

1. Basyouni R, Parkinson C (2022): Mapping the social landscape: tracking patterns of interpersonal relationships. Trends in Cognitive Sciences 26: 204–221.

2. Ciranka S, Lin Y, Wittmann M (2025): Basis functions for social decision-making develop during adolescence.

3. Lin Y, Pellicano E, Dickson C, Trudel N, Noonan M, Lockwood P, et al. (2026): Preserved self-other integration during social decision making among individuals with elevated autistic traits. bioRxiv 2026–06.

4. Wittmann MK, Lin Y, Pan D, Braun MN, Dickson C, Spiering L, et al. (2025): Basis functions for complex social decisions in dorsomedial frontal cortex. Nature. 10.1038/s41586-025-08705-9

5. Beeney JE, Hallquist MN, Ellison WD, Levy KN (2016): Self–other disturbance in borderline personality disorder: Neural, self-report, and performance-based evidence. Personality Disorders: Theory, Research, and Treatment 7: 28–39.

6. Blatt SJ, Luyten P (2009): A structural-developmental psychodynamic approach to psychopathology: two polarities of experience across the life span. Dev Psychopathol 21: 793–814.

7. De Meulemeester C, Luyten P, Fonagy P (2025): Special section: Self–other distinction in personality disorders. Personality Disorders: Theory, Research, and Treatment 16: 103–109.

8. Korn CW, La Rosée L, Heekeren HR, Roepke S (2016): Social feedback processing in borderline personality disorder. Psychol Med 46: 575–587.

9. Jørgensen CR, Bøye R (2022): How Does It Feel to Have a Disturbed Identity? The Phenomenology of Identity Diffusion in Patients With Borderline Personality Disorder: A Qualitative Study. Journal of Personality Disorders 36: 40–69.

10. De Meulemeester C, Lowyck B, Vermote R, Verhaest Y, Luyten P (2017): Mentalizing and interpersonal problems in borderline personality disorder: The mediating role of identity diffusion. Psychiatry Research 258: 141–144.

11. Sollberger D, Gremaud-Heitz D, Riemenschneider A, Küchenhoff J, Dammann G, Walter M (2012): Associations between Identity Diffusion, Axis II Disorder, and Psychopathology in Inpatients with Borderline Personality Disorder. Psychopathology 45: 15–21.

12. Fonagy P, Luyten P (2009): A developmental, mentalization-based approach to the understanding and treatment of borderline personality disorder. Dev Psychopathol 21: 1355–1381.

13. Liebke L, Bungert M, Thome J, Hauschild S, Gescher DM, Schmahl C, et al. (2017): Loneliness, social networks, and social functioning in borderline personality disorder. Personality Disorders: Theory, Research, and Treatment 8: 349–356.

14. Reinhard MA, Nenov-Matt T, Padberg F (2022): Loneliness in Personality Disorders. Curr Psychiatry Rep 24: 603–612.

15. Schulze A, Streit F, Zillich L, Awasthi S, Hall ASM, Jungkunz M, et al. (2023): Evidence for a shared genetic contribution to loneliness and borderline personality disorder. Transl Psychiatry 13: 398.

16. Wicher CL, Dombrovski AY, Hallquist MN, Buecker S, Wright AG, Kaurin A (2025): Daily loneliness and suicidal ideation in borderline personality disorder. Personality Disorders: Theory, Research, and Treatment.

17. Mermin SA, Steigerwald G, Choi-Kain LW (2025): Borderline Personality Disorder and Loneliness: Broadening the Scope of Treatment for Social Rehabilitation. Harvard Review of Psychiatry 33: 31.

18. Vonderlin R, Claus C, Hanraths S, Lerchl AS, Senyüz B, Kleindienst N, et al. (2026): Loneliness in borderline personality disorder: The role of misalignments between self-view and social expectations in social value orientation and justice sensitivity. Comprehensive Psychiatry 146: 152663.

19. Barnby JM, Nguyen J, Griem J, Wloszek M, Burgess H, Richards LJ, et al. (2025): Self-other generalisation shapes social interaction and is disrupted in borderline personality disorder. eLife 14: RP104008.

20. Gregorova K, Waltmann M, Will G-J, Mittermeier S, Bertsch K, Romanos M, et al. (n.d.): Young individuals with Borderline Personality Disorder do not learn they are liked and show altered self-esteem reactivity to social feedback.

21. Niedtfeld I (2017): Experimental investigation of cognitive and affective empathy in borderline personality disorder: Effects of ambiguity in multimodal social information processing. Psychiatry Res 253: 58–63.

22. Herpertz SC, Bertsch K (2014): The social-cognitive basis of personality disorders. Current Opinion in Psychiatry 27: 73.

23. Story GW, Ereira S, Valle S, Chamberlain SR, Grant JE, Dolan RJ (2024): A computational signature of self-other mergence in Borderline Personality Disorder. Transl Psychiatry 14: 473.

24. Meulemeester CD (2021): The role of impairments in self–other distinction in borderline personality disorder: A narrative review of recent evidence. Neuroscience and Biobehavioral Reviews 13.

25. King-Casas B, Sharp C, Lomax-Bream L, Lohrenz T, Fonagy P, Montague PR (2008): The Rupture and Repair of Cooperation in Borderline Personality Disorder. Science 321: 806–810.

26. Chang L, Tsao DY (2017): The Code for Facial Identity in the Primate Brain. Cell 169: 1013–1028 e14.

27. Hunt JJ, Dayan P, Goodhill GJ (2013): Sparse Coding Can Predict Primary Visual Cortex Receptive Field Changes Induced by Abnormal Visual Input ((M. Bethge, editor)). PLoS Comput Biol 9: e1003005.

28. Ereira S, Hauser TU, Moran R, Story GW, Dolan RJ, Kurth-Nelson Z (2020): Social training reconfigures prediction errors to shape Self-Other boundaries. Nat Commun 11: 3030.

29. Pronizius E, Bukowski H, Lamm C (2024): Comparing self–other distinction across motor, cognitive and affective domains. R Soc Open Sci 11: 240662.

30. Wittmann MK, Trudel N, Trier HA, Klein-Flügge MC, Sel A, Verhagen L, Rushworth MFS (2021): Causal manipulation of self-other mergence in the dorsomedial prefrontal cortex. Neuron 109: 2353–2361.e11.

31. Wittmann MK, Kolling N, Faber NS, Scholl J, Nelissen N, Rushworth MF (2016): Self-Other Mergence in the Frontal Cortex during Cooperation and Competition. Neuron 91: 482–93.

32. Chierchia G, Fuhrmann D, Knoll LJ, Pi-Sunyer BP, Sakhardande AL, Blakemore S-J (2019): The matrix reasoning item bank (MaRs-IB): novel, open-access abstract reasoning items for adolescents and adults. Royal Society Open Science 6: 190232.

33. Roitman JD, Shadlen MN (2002): Response of neurons in the lateral intraparietal area during a combined visual discrimination reaction time task. The Journal of neuroscience : the official journal of the Society for Neuroscience 22: 9475–89.

34. Lockwood PL, Wittmann MK, Apps MA, Klein-Flügge MC, Crockett MJ, Humphreys GW, Rushworth MF (2018): Neural mechanisms for learning self and other ownership. Nature communications 9: 4747.

35. Sui J, Gu X (2017): Self as Object: Emerging Trends in Self Research. Trends Neurosci. 10.1016/j.tins.2017.09.002

36. Neuhaus JM, Kalbfleisch JD (1998): Between- and Within-Cluster Covariate Effects in the Analysis of Clustered Data. Biometrics 54: 638.

37. Kesebir S, Oishi S (2010): A Spontaneous Self-Reference Effect in Memory: Why Some Birthdays Are Harder to Remember Than Others. Psychol Sci 21: 1525–1531.

38. Yin S, Sui J, Chiu Y-C, Chen A, Egner T (2019): Automatic Prioritization of Self-Referential Stimuli in Working Memory. Psychol Sci 30: 415–423.

39. Humphreys GW, Sui J (2015): The salient self: Social saliency effects based on self-bias. Journal of Cognitive Psychology 27: 129–140.

40. Müller-Pinzler L, Czekalla N, Mayer AV, Schröder A, Stolz DS, Paulus FM, Krach S (2022): Neurocomputational mechanisms of affected beliefs. Commun Biol 5: 1241.

41. Schröder A, Czekalla N, Mayer AV, Zhang L, Stolz DS, Korn CW, et al. (2024, August 30): Computational Modeling shows Confirmation Bias during Formation and Revision of Self-Beliefs. 10.1101/2024.08.30.610443

42. Sui J, Humphreys GW (2015): The Integrative Self: How Self-Reference Integrates Perception and Memory. Trends in Cognitive Sciences 19: 719–728.

43. Miyamoto K, Harbison C, Tanaka S, Saito M, Luo S, Matsui S, et al. (2025): Asymmetric projection of introspection reveals a behavioural and neural mechanism for interindividual social coordination. Nat Commun 16: 295.

44. American Psychiatric Association (2022): *Diagnostic and Statistical Manual of Mental Disorders*, 5th, text revision ed. American Psychiatric Association Publishing. 10.1176/appi.books.9780890425787

45. World Health Organization (2019): International Statistical Classification of Diseases and Related Health Problems (11th *Ed.)*. World Health Organization. Retrieved from https://icd.who.int/

46. Korn CW, La Rosée L, Heekeren HR, Roepke S (2016): Processing of information about future life events in borderline personality disorder. Psychiatry Research 246: 719–724.

47. Jeung H, Walther S, Korn CW, Bertsch K, Herpertz SC (2018): Emotional responses to receiving peer feedback on opinions in borderline personality disorder. Personality Disorders: Theory, Research, and Treatment 9: 595–600.

48. Doppelhofer LM, Perla R, Frolichs KMM, Rosenblau G, Herpertz SC, Korn CW (2026): Estimating and learning personality traits of and from women with borderline personality disorder. bord personal disord emot dysregul 13: 5.

49. Bender DS, Skodol AE (2007): Borderline personality as a self-other representational disturbance. Journal of personality disorders 21: 500–517.

50. Hanegraaf L, Hohwy J, Verdejo-Garcia A (2022): Latent classes of maladaptive personality traits exhibit differences in social processing. Journal of Personality 90: 615–630.

51. Allport FH (1924): The group fallacy in relation to social science. Journal of Abnormal and social Psychology 19: 60–73.

52. Barrett LF, Miller EK (2026): Categorization is ‘baked’ into the brain. Nat Rev Neurosci. 10.1038/s41583-026-01036-2

53. De Dreu CKW, Gross J, Fariña A, Ma Y (2020): Group Cooperation, Carrying-Capacity Stress, and Intergroup Conflict. Trends in Cognitive Sciences 24: 760–776.

54. Hein G, Silani G, Preuschoff K, Batson CD, Singer T (2010): Neural responses to ingroup and outgroup members’ suffering predict individual differences in costly helping. Neuron 68: 149–60.

55. Paus T, Keshavan M, Giedd JN (2008): Why do many psychiatric disorders emerge during adolescence? Nat Rev Neurosci 9: 947–957.

56. Solmi M, Radua J, Olivola M, Croce E, Soardo L, Salazar De Pablo G, et al. (2022): Age at onset of mental disorders worldwide: large-scale meta-analysis of 192 epidemiological studies. Mol Psychiatry 27: 281–295.

57. Hanegraaf L, Van Baal S, Hohwy J, Verdejo-Garcia A (2021): A systematic review and meta-analysis of ‘Systems for Social Processes’ in borderline personality and substance use disorders. Neuroscience & Biobehavioral Reviews 127: 572–592.

58. De Panfilis C, Riva P, Preti E, Cabrino C, Marchesi C (2015): When social inclusion is not enough: Implicit expectations of extreme inclusion in borderline personality disorder. Personality Disorders: Theory, Research, and Treatment 6: 301–309.

59. Gerra ML, Ardizzi M, Martorana S, Leoni V, Riva P, Preti E, et al. (2021): Autonomic vulnerability to biased perception of social inclusion in borderline personality disorder. bord personal disord emot dysregul 8: 28.

60. Bungert M, Liebke L, Thome J, Haeussler K, Bohus M, Lis S (2015): Rejection sensitivity and symptom severity in patients with borderline personality disorder: effects of childhood maltreatment and self-esteem. bord personal disord emot dysregul 2: 4.

61. Cavicchioli M, Maffei C (2020): Rejection sensitivity in borderline personality disorder and the cognitive–affective personality system: A meta-analytic review. Personality Disorders: Theory, Research, and Treatment 11: 1–12.

62. Gutz L, Renneberg B, Roepke S, Niedeggen M (2015): Neural processing of social participation in borderline personality disorder and social anxiety disorder. Journal of Abnormal Psychology 124: 421–431.

63. Weinbrecht A, Niedeggen M, Roepke S, Renneberg B (2018): Feeling excluded no matter what? Bias in the processing of social participation in borderline personality disorder. NeuroImage: Clinical 19: 343–350.

64. Bateman A, Fonagy P, Allen JG (2009): Theory and practice of mentalization based therapy. Retrieved from https://api.semanticscholar.org/CorpusID:148096114

65. Nolte T, Hutsebaut J, Sharp C, Campbell C, Fonagy P, Bateman A (2023): The role of epistemic trust in mentalization-based treatment of borderline psychopathology. Journal of Personality Disorders 37: 633–659.

66. Linehan M (1993): Cognitive-Behavioral Treatment of Borderline Personality Disorder. Guilford press.

67. Baek EC, Hyon R, López K, Du M, Porter MA, Parkinson C (2023): Lonely Individuals Process the World in Idiosyncratic Ways. Psychol Sci 34: 683–695.

68. Courtney AL, Meyer ML (2020): Self-Other Representation in the Social Brain Reflects Social Connection. J Neurosci 40: 5616–5627.

69. Hawkley LC, Cacioppo JT (2010): Loneliness Matters: A Theoretical and Empirical Review of Consequences and Mechanisms. ann behav med 40: 218–227.

70. Euler S, Nolte T, Constantinou M, Griem J, Montague PR, Fonagy P (2021): Personality and Mood Disorders Research Network.(2021). Interpersonal problems in borderline personality disorder: Associations with mentalizing, emotion regulation, and impulsiveness. Journal of personality disorders 35: 177–193.

71. Lieb K, Zanarini MC, Schmahl C, Linehan MM, Bohus M (2004): Borderline personality disorder. The lancet 364: 453–461.

72. Nenov-Matt T, Barton BB, Dewald-Kaufmann J, Goerigk S, Rek S, Zentz K, et al. (2020): Loneliness, Social Isolation and Their Difference: A Cross-Diagnostic Study in Persistent Depressive Disorder and Borderline Personality Disorder. Front Psychiatry 11: 608476.

73. Haihambo N, Layiwola D, Blank H, Hurlemann R, Scheele D (2025): Loneliness and social conformity: A predictive processing perspective. Ann NY Acad Sci 1547: 5–17.

74. Mittermeier S, Gregorova K, Goettfert C, Merz C, Weiß M, Krauss J, et al. (2025): aVeRsive tension: A new virtual reality paradigm to assess emotional arousal in adolescent and young adult patients with symptoms of borderline personality disorder. International Journal of Clinical and Health Psychology 25: 100583.

75. Gunderson JG, Herpertz SC, Skodol AE, Torgersen S, Zanarini MC (2018): Borderline personality disorder. Nat Rev Dis Primers 4: 18029.

76. Gawda B, Czubak K (2017): Prevalence of Personality Disorders in a General Population Among Men and Women. Psychol Rep 120: 503–519.

77. Ten Have M, Verheul R, Kaasenbrood A, Van Dorsselaer S, Tuithof M, Kleinjan M, De Graaf R (2016): Prevalence rates of borderline personality disorder symptoms: a study based on the Netherlands Mental Health Survey and Incidence Study-2. BMC Psychiatry 16: 249.

78. Fleischhaker C, Sixt B, Schulz E (2011): DBT-A Dialektisch-behaviorale Therapie für Jugendliche: Ein Therapiemanual mit Arbeitsbuch auf CD. Berlin, Heidelberg: Springer Berlin Heidelberg. 10.1007/978-3-642-13008-3

79. Chanen AM, Kaess M (2012): Developmental pathways to borderline personality disorder. Current psychiatry reports 14: 45–53.

80. Kaess M, Fischer-Waldschmidt G, Resch F, Koenig J (2017): Health related quality of life and psychopathological distress in risk taking and self-harming adolescents with full-syndrome, subthreshold and without borderline personality disorder: rethinking the clinical cut-off? Borderline personality disorder and emotion dysregulation 4: 7.

81. BSCL: Brief-Symptom-Checklist: Manual (n.d.):

82. De Leeuw JR (2015): jsPsych: A JavaScript library for creating behavioral experiments in a Web browser. Behav Res 47: 1–12.

83. Inc TM (2023): MATLAB version: 14.6.1 (R2023b). Natick, Massachusetts, United States: The MathWorks Inc. Retrieved from https://www.mathworks.com

84. JASP Team (2025): JASP (Version 0.19.3)[Computer software]. Retrieved from https://jasp-stats.org/

85. Wittmann MK, Fouragnan E, Folloni D, Klein-Flügge MC, Chau BKH, Khamassi M, Rushworth MFS (2020): Global reward state affects learning and activity in raphe nucleus and anterior insula in monkeys. Nat Commun 11: 3771.

86. Huth AG, de Heer WA, Griffiths TL, Theunissen FE, Gallant JL (2016): Natural speech reveals the semantic maps that tile human cerebral cortex. Nature 532: 453–8.

87. Bates D, Mächler M, Bolker B, Walker S (2015): Fitting Linear Mixed-Effects Models Using lme4. Journal of Statistical Software 67: 1–48.

88. Kuznetsova A, Brockhoff PB, Christensen RHB (2017): lmerTest Package: Tests in Linear Mixed Effects Models. J Stat Soft 82. 10.18637/jss.v082.i13

89. Lenth R, Singmann H, Love J, Buerkner P, Herve M (2019): Package “emmeans.”(R Package, Version, 1 3.2)[Computer software].

